# The role of *swnR* gene on the biosynthesis pathway of the swainsonine in *Metarhizium anisopliae*

**DOI:** 10.1101/813295

**Authors:** Lu Sun, Runjie Song, Jinglong Wang, Yiling Liu, Yu Zhang, Yanli Zhu, Qingyun Guo, Chonghui Mo, Baohai Wang, Baoyu Zhao, Hao Lu

**Author notes:** Corresponding author. Lu H. These authors contributed equally to this work (Lu Sun, Runjie Song, Jinglong Wang).

## Abstract

Swainsonine (SW) is the principal toxic ingredient of locoweeds, and is produced by fungi including *Metarhizium anisopliae*, *Slafractonia leguminicola*, and *Alternaria oxytropis*. Studies of the SW biosynthesis pathway in these fungi have demonstrated the requirement for a *swn*K gene and the presence of a variety of other SWN cluster genes, but have not determined a precise role for the *swnR* gene, which encodes a NADB Rossmann-fold reductase, nor if it is necessary for the biosynthesis of SW. In this study, we used homologous recombination (HR) to knock out the *swnR* gene of *M. anisopliae* to determine its effect on the SW biosynthesis pathway. The concentration of SW was measured in the fermentation broth of *M. anisopliae* at 1 d, 3 d, 5 d and 7 d using a Q Exactive Mass Spectrometer. The gene for swnR was detected by RT-qPCR. To determine the role of the *swnR* gene in the SW biosynthesis pathway of *M. anisopliae*, we used PEG-mediated homologous recombination (HR) to transform a wild-type strain (WT) with a Benomyl (*ben*)-resistant fragment to knock out the *swnR* gene producing a mutant-type strain (MT). A complemented-type (CT) strain was produced by adding a complementation vector that contains the glufosinate (herbicide) resistance (*bar*) gene as a marker. The content of SW decreased, but was not eliminated in the fermentation broth of the MT strain, and returned to the original level in the CT strain. These results indicate that the *swnR* gene plays a crucial role in the SW biosynthesis pathway of *M. anisopliae*, but suggests that another gene in the fungus may share the function of swnR.

## Introduction

*Metarhizium anisopliae* is anentomopathogenic fungus that can infect a variety of agricultural pest insects and produces swainsonine[1–3]. Swainsonine is an indolizidine alkaloid that inhibits alpha-mannosidases. Swainsonine is produced by a variety of fungi and is the major toxic component of locoweed plants, which are widely found in western China and North America [4–9]. Swainsonine (SW) causes locoism in grazing animals [10–12], when consumed for an extended time. SW also has anti-tumor properties and may enhance the body’s immunity [13–16].

Recently, Cook et al. [17] sequenced the genome of two swainsonine-producing fungi *Slafractonia leguminicola* and *Alternaria oxytropis*, and compared their swainsonine producing genes to those in *M. robertsii*and other swainsonine-producing fungi. All the swainsonine-producing fungi contained a common gene cluster "*SWN*", which included *swnH_1_*, *swnH_2_*, *swnK*, *swnN*, *swnR*, *swnA*, and *swnT*. Fungi that do not produce swainsonine do not contain the SWN cluster. These genes encode catalytic enzymes involved in the SW biosynthesis. *swnN* and *swnR* both encode NADB Rossmann-fold reductases. After annotating the "*SWN*" gene cluster, Cook et al. [17] successfully constructed a *swnK* knockout vector for the entomopathogen *M. robertsii*, and obtained a *swnK* mutant strain by HR. The LC-MS analysis showed that SW was eliminated in the fermentation broth of the mutant strain and the content of SW returned to normal after complementation with the *swnK* gene, indicating that the *swnK* gene is essential for SW biosynthesis.

The precise roles for the other genes in the SWN cluster were not determined. The aims of this study were to investigate the role of the *swnR* gene in the SW biosynthesis pathway of *M. anisopliae*. To accomplish this, *swnR* of *M. anisopliae* was knocked out using HR. The deletion of *swnR* led to a significant decrease in the SW content in the fermentation broth of *M. anisopliae*. The content of SW in the complemented strain was similar to that found for the wild type strain.

## Materials and methods

### Strain and fermentation culture

*Metarhizium anisopliae*, obtained from Xi’an Jin Berry Biological Technology Co. Ltd., China, was inoculated onto Sabouraud medium (SDA) [18] containing 50 µg/ml chloramphenicol and cultured at 30 °C for 10 days. Six 5 mm plugs of mycelium were transferred into 200 mL SDA liquid culture media (without agar) and fermented at 28 °C, 180 rpm, for 1 d, 3 d, 5 d, or 7 d. The mycelium and the fermentation broth were filtered through three layers of Miracloth (EMD Millipore Corp, Billerica, MA, USA) to collect the mycelium. No conidia were detected from the cultures. The mycelium was dried at room temperature and stored at 4 °C for use.

### RT-qPCR analysis of genes in the SW biosynthesis pathway of *M. anisopliae*

Fungal RNA was extracted using the E.Z.N.A. Fungal RNA Kit (Omega). RNA was reverse transcribed into DNA by using PrimeScript™RT reagent Kit (Takara). Primers for amplification of SwnN, SwnT, SwnK, SwnH1, SwnH2, SwnR, and 18SrRNA are shown in Table S1. All of the genes were amplified using RT-qPCR with the following conditions: 1 cycle of 95 °C for 10 min; 40 cycles of denaturation at 95 °C for 10 s, annealing at 55 °C for 90 s, and an extension at 72 °C for 32 s. Cultures were tested using RT-qPCR at 1d, 3d, 5d, and 7d.

### Identification of swnR gene of *M. anisopliae*

Fungal DNA was extracted using the CTAB method. Primers for amplification of *swnR* (Table S2) were designed from the *swnR* sequence from Cook et al. [17], GeneBankKID61009 of *M. anisopliae* ARSEF 549. The *swnR* gene was amplified from *M. anisopliae* DNA using L1/R1 primers (Text S1).

### Vector construction

The upstream and downstream fragments of the *swnR* gene (Fig.S1) and the benomyl (fungicide) resistance gene (*ben*) (Fig.S2) were inserted into pUC19 (Takara) digested with *Eco*R I/*Bam*H I (Takara) (Fig.S3) using the In-Fusion® HD Cloning System (Takara) to construct a knockout construct targeting the *swnR* gene (Fig.S4, Fig. S5 andText S1).

The primers L3 and R3 (Table S1) were used to amplify the *ben* resistance gene from pBARGPE1-BenA (Wuhan Jingxiu Scientific Biotechnology Co., Ltd., China) as a template. The primers L2/R2 and L4/R4 (Table S1) were used to amplify the upstream target fragment (sw*nR*-I) and the downstream target fragment (*swnR*-II), respectively, of the *swnR* gene from the genomic DNA of *M. anisopliae*. The *swnR*-I, *ben*, *swnR*-II and the double-cut pUC19 vector were ligated using In-Fusion cloning. The *swnR* gene fragment was amplified using primers L1/R1 (Table S1) from the genomic DNA of *M. anisopliae*. To produce a complementation vector, the *swnR* was inserted between *trpC* promoter and *trpC* terminator of pBARGPE1 vector, which contains the glufosinate (herbicide) resistance (*bar*) gene as a marker, using In-Fusion cloning (Fig.1).

**Fig. 1.**
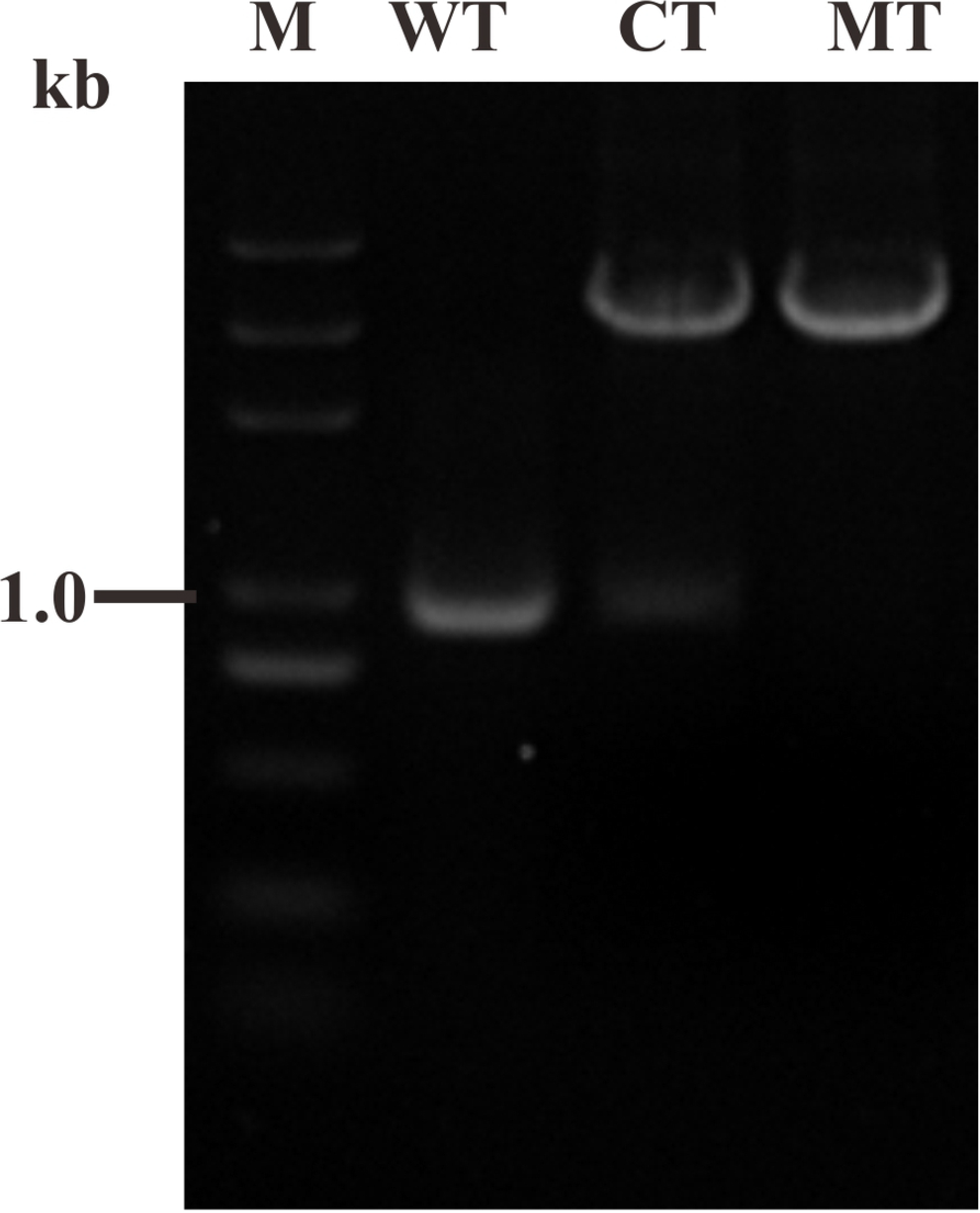
Verification of MT, CT and WT. PCR products using primers L1/R1 from WT (wild type), knockout (MT), and complement (CT).

### Preparation of protoplasts

Six 5 mm plugs from 10-day cultures of *M. anisopliae* grown on SDA media were transferred into each 200 mL flask of SDA liquid culture media (without agar), and incubated at 28 °C, 180 rpm for 1 d, 2 d, 3 d, 4 d and 5 d. The resulting mycelia were filtered through sterile miracloth. To the collected hyphae were added different concentrations of enzymatic hydrolysate (Sigma Aldrich) prepared with 1.2 M KCl, and hydrolyzed at 30 °C, 100 rpm, for 1 h, 3 h, 5 h, 7 h, and 10 h. The optimal combination of enzymes and conditions were determined based on protoplast yield. Yield from different enzymes, including 1% snail enzyme, 1% cellulase, and 1% lysing enzymes, and combinations of the enzymes were also tested. The enzymatically digested mixtures were filtered through a layer of sterile miracloth and two layers of filter paper into a sterile 50 mL centrifuge tube, and the protoplasts were washed extensively with 1.2 M KCl and centrifuged at 4000 rpm for 6 min at room temperature. After discarding the supernatant, 10 mL of STC Buffer (0.6 M Sorbitol; 10 mM Tris-HCl; 10 mM CaCl_2_, pH 6.5) was added and the protoplasts were gently resuspended. The mixture was centrifuged at 4000 rpm for 6 minutes. After discarding the supernatant, 1 mL of STC Buffer was added. The protoplasts were then centrifuged at room temperature at 3500 rpm for 6 min, which was repeated. Finally, protoplasts were adjusted to 2-5×10^7^/mL for subsequent experiments.

### PEG mediated DNA transformation

Transformation of the protoplasts were done as in Proctor et al., 1995[19]. Approximately 5-10 µg of the linearized *swnR* knockout vector was added to a 50 mL centrifuge tube containing 2-5×10^7^/mL protoplasts, and allowed to stand at room temperature for 20 min without shaking. Then 1-1.25 mL of 40% PTC (40% PEG 8000, 20% sucrose, 50 mM CaCl_2_, 10 mM Tris-HCl) was added to the tube (mixed thoroughly by inversion), and let stand at room temperature for 20 min without shaking. Thereafter, 5 mL of TB_3_ (0.3% Yeast Extract, 0.3% acid hydrolyzed casein, 20% sucrose) containing 50 g/mL ampicillin (Sigma Aldrich) was added and shaken at room temperature overnight. The overnight protoplasts were centrifuged at 4000 rpm for 6 min, the supernatant was discarded, and about 1 mL of the remaining liquid was used to suspend the remainder. The regenerated protoplasts were added to 10 mL of Bottom Agar (0.3% Yeast Extract, 0.3% acid hydrolyzed casein, 20% sucrose, 1% Agar) containing 100 µg/mL benomyl (Fig.S4). After incubation at 30 °C for 10 hours, Top Agar (0.3% Yeast Extract, 0.3% acid hydrolyzed casein, 20% sucrose, 1.5% Agar) containing 200 µg/mL benomyl was added. After 3-5 d, a single colony transformant grew on the plate, which was transferred to SDA medium containing 200 µg/mL benomyl. The wild type *M. anisopliae* was used as a control. The *swnR* gene mutant strain of *M. anisopliae* was named MT. The transformation of the complement vector was the same as described above, and 2 mg/mL of glufosinate (Fig.S6) was used for screening of the complement (CT).

### PCR identification of MT and CT

The *M. anisopliae* strain carrying the benomyl resistance gene was used as a template, and PCR amplification was carried out using primer L1/R1. Subsequently, the complemented strain was subjected to the same PCR amplification using primers L1/R1 (Table S1).

### Phenotypic observation and growth rate determination MT, CT and WT

Colonies of MT and WT of the same size were inoculated into the same position on the SDA medium and grown at 28 °C for 3 d, 5 d and 10 d, after which they were measured for diameter and photographed.

### SW content detection of fermentation broth of WT, MT, CT in *M. anisopliae*

The WT, MT, CT strains were inoculated into SDA medium containing 50 µg/ml chloramphenicol and cultured at 28 °C for 10 days. Then six 5 mm plugs of each strain were transferred into 200 mL flasks of SDA (without agar) culture medium and grown at 28 °C, 180 rpm, for 3 d. The flasks of fermentation broth of WT, MT, and CT were combined by strain and filtered to obtain 500 mL of fermentation broth. The SW (control from Sigma Aldrich) in each extract was analyzed using Q Exactive Mass Spectrometer (Thermo Fisher) using the methods of Song et al. [20]. SW concentration was tested three times for each strain.

### Statistical analysis

In this study, each measurement was tested three times. Statistical analysis was performed on the measured data using SPSS 20.0 software. The results were expressed as mean ± SEM. One-way ANOVA was performed on each sample, *P<0.05, indicating a significant difference between the two groups, **P<0.01, indicating that the difference between the two groups is highly significant. Results from the cultures were used for determining the optimal time periods for swainsonine production. The mass concentration peak area for SW was compared using linear regression. The colony diameters were measured by ruler. RT-qPCT data were analyzed using the 2 ^−△△^CT method.

## Results

### Detection of SW in fermentation broth of *M. anisopliae*in different periods

To detect the SW content in the fermentation broth at different time points, the same volume of *M. anisopliae* fermentation broth was concentrated, and the level of SW was detected using a Q Exactive Mass Spectrometer. The retention times for the SW peak of the SW standard and the SW test samples was 4.4 (Fig.S7A, S7B,S7C, S7D, and S7E). The standard curve was drawn according to the calculated regression equation: Y=31302.5X-45910.5 (R^2^= 0.997) (Fig.S7F). From the linear regression equation of mass concentration-peak area of SW, the SW content in the fermentation broth of *M. anisopliae* was calculated to be 169.67 ± 50.78 μg/mg (S7A), 174.01 ± 45.79 μg/mg (Fig.S7B), 116.72 ± 45.74 μg/mg (Fig.S7C), and 104.85 ± 40.35 μg/mg (Fig. S7D), at 1, 3, 5, and 7 days of growth, respectively. Day 3 of fermentation gave the highest content of SW (Fig. 2).

**Fig. 2.**
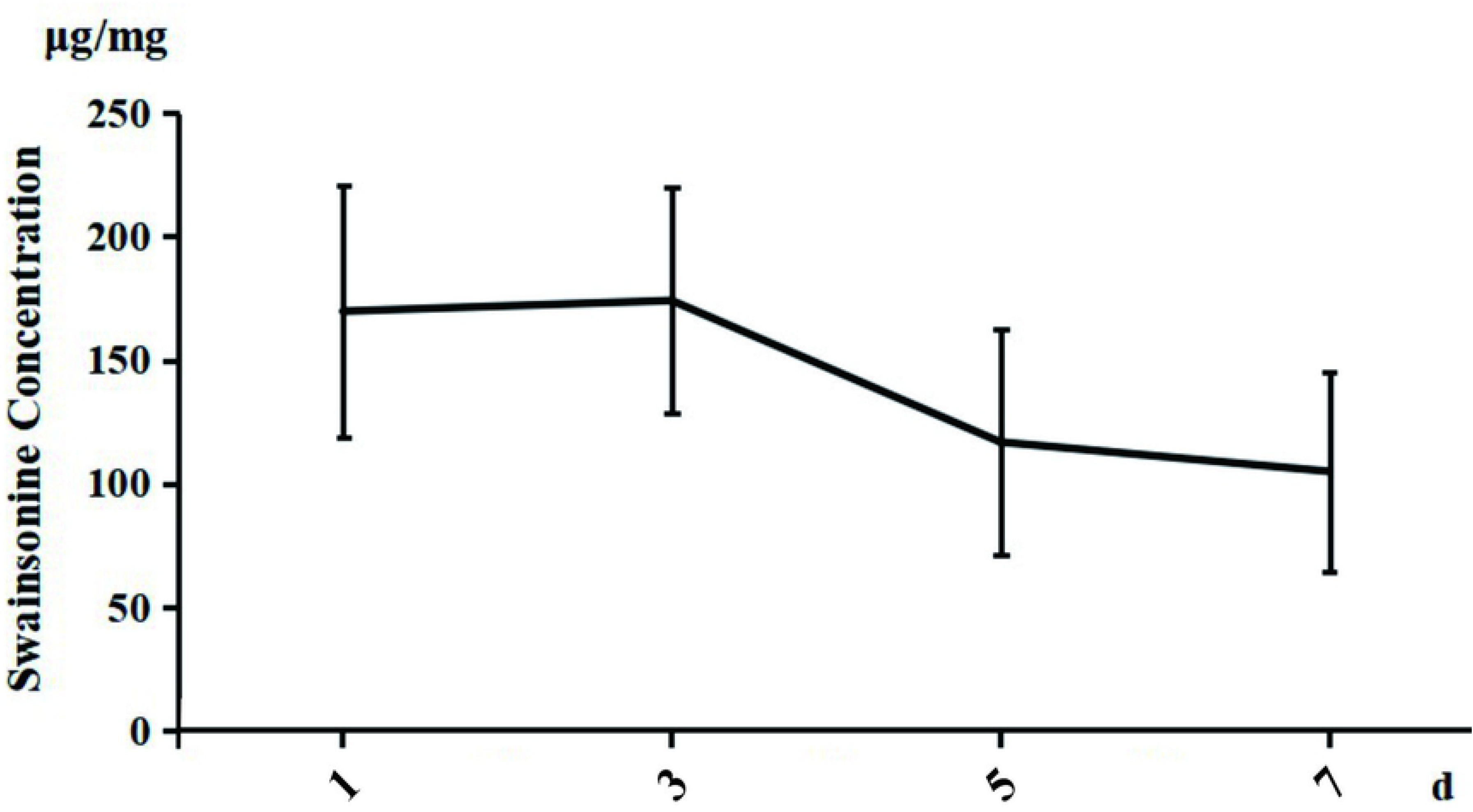
Detection of SW in fermentation broth of *M. anisopliae* in different time. Time course - SW amount graph, There are three parallel replicates for each test sample in this test. Each data in the graph is the mean ± SEM, n = 3, P<0.05.

### RT-qPCR analysis of key catalytic enzyme genes in the SW biosynthesis pathway of *M. anisopliae*

RT-qPCR was conducted for genes in the SW biosynthesis pathway to determine their relative expression at 1d, 3d, 5d, and 7d. The expression of *swnN* was down-regulated from 1d to 3d. The expression levels of *swnT* and *swnK* decreased at 3 d and 5 d, and the expression of *swnH*_*1*_ and *swnH*_*2*_ did not significantly change. The expression of *swnR* gene was significantly up-regulated at 3 d and decreased at 5d and 7d (Fig.3).

**Fig. 3.**
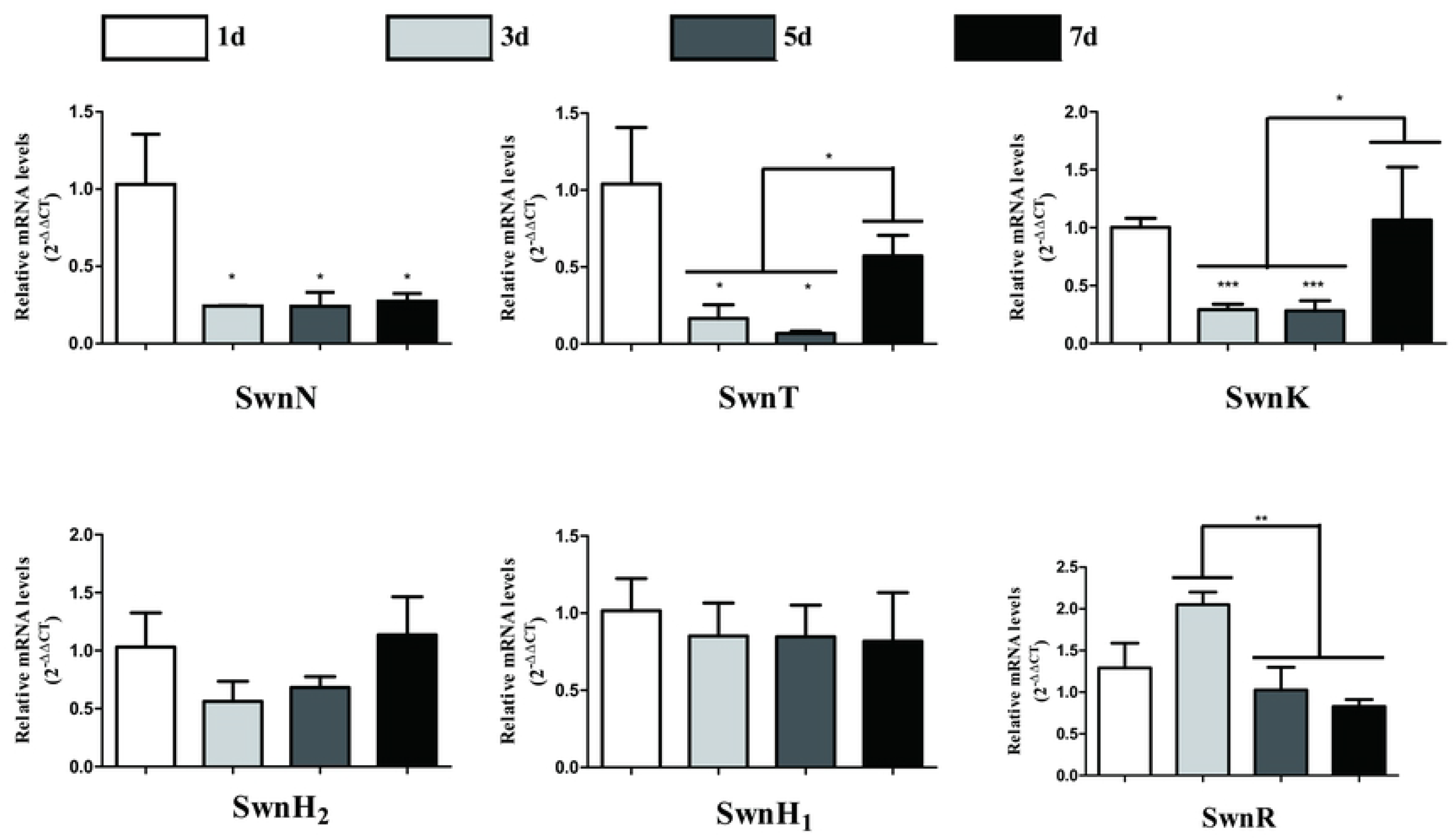
RT-qPCR analysis of key catalytic enzyme genes in the SW biosynthesis pathway of *M. anisopliae*. RNA was extracted, converted to cDNA, and the expression of *swnN*, *swnT, swnK*, *swnH*_*2*_, *swnH*_*1*_, *swnA* and *swnR* in *M. anisopliae*at 1 d, 3 d, 5 d and 7 d was tested. Error bars represent the standard error of the mean (n = 3), *P< 0.05; **P<0.01.

### Screening for optimal conditions for protoplast preparation

We explored the conditions affecting the preparation of protoplasts of *M. anisopliae*to enhance genetic manipulation of the fungus. Hyphae produced from fermentation times of 1 d, 2 d, 3 d, 4 d, and 5 d were enzymatically digested for 3 h at 30 °C, 100 rpm. The optimal fermentation time of 2 d was determined based on protoplast yield (Fig.4A). The effects of different enzymatic hydrolysis combinations and enzymatic concentrations were compared on protoplast preparations of *M. anisopliae*. Enzymatic hydrolysis using 1% snail enzyme, 1% cellulase, and 1% lysing enzymeswas the best and produced the largest number of protoplasts (Fig.4B). Duration of enzymatic hydrolysis time is also an important variable of the protoplast preparation process, so the optimal enzymatic hydrolysis time was assessed. When enzymatically hydrolyzed for 3 h, the hyphal wall was completely dissolved, releasing large numbers of protoplasts (Fig.4C and 4D).

**Fig. 4.**
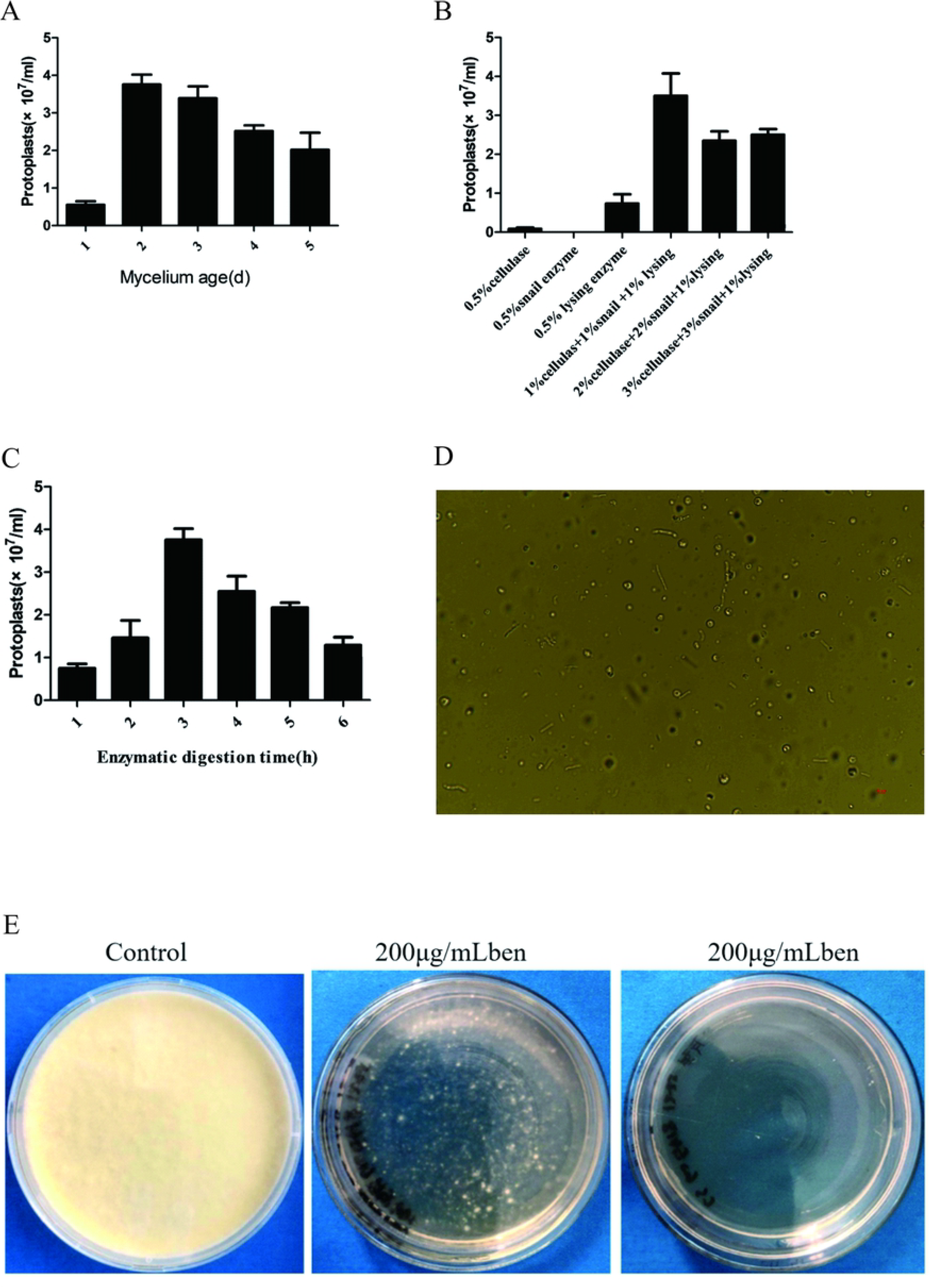
Screening for optimal conditions for protoplast preparation and PEG mediated *swnR* transformation. (A-D) *M. anisopliae* cultured for 10 d were inoculated to 200 mL of Czapek’s medium with a sterile inoculation needle, cultured at 30 °C, 180 rpm for 1 d, 2 d, 3 d, 4 d and 5 d. The mycelium of the above fermentation for 1 d, 2 d, 3 d, 4 d and 5 d were filtered with sterile miracloth. The collected hyphae were added to different concentrations of enzymatic hydrolysate prepared with 1.2 M KCl as osmotic stabilizer, and hydrolyzed at 30 °C, 100 rpm, for 1 h, 3 h, 5 h, 7 h and 10 h. The protoplast release of *M. anisopliae* was the highest when digested with 1% snail enzyme, 1% cellulase and 1% lysing enzymes for 3 h. Each data in the graph is the mean ± SEM, n = 3, P<0.05.

### Production of MT and CT

To determine the role of the *swnR* gene in the SW biosynthesis pathway of *M. anisopliae*, homologous recombination was used to knock out *swnR.* The resulting transformant (1 out of 55) grew on SDA media containing 200 μg/mL benomyl. Subsequently, the L1/R1 primer set was used to identify the genomic DNA of the transformant (MT) using electrophoresis and sequencing (Fig.1 andText S2). To verify the status of MT, a complement was produced by transforming the wild-type *swnR* gene in pBARGPE1 into the MT. The complement transformant was grown on SDA medium containing 2 mg/mL glufosinate and was identified as above.

### Phenotypic observation and growth rate determination of WT, MT, and CT

TheWT, MT, and CT isolates were grown on three SDA plates each with two colonies per plate for 3 d, 5 d, and 10 d (Fig.5A) to compare growth. The colony diameters, phenotypes, and growth rates did not change significantly (Fig.5B).

**Fig. 5.**
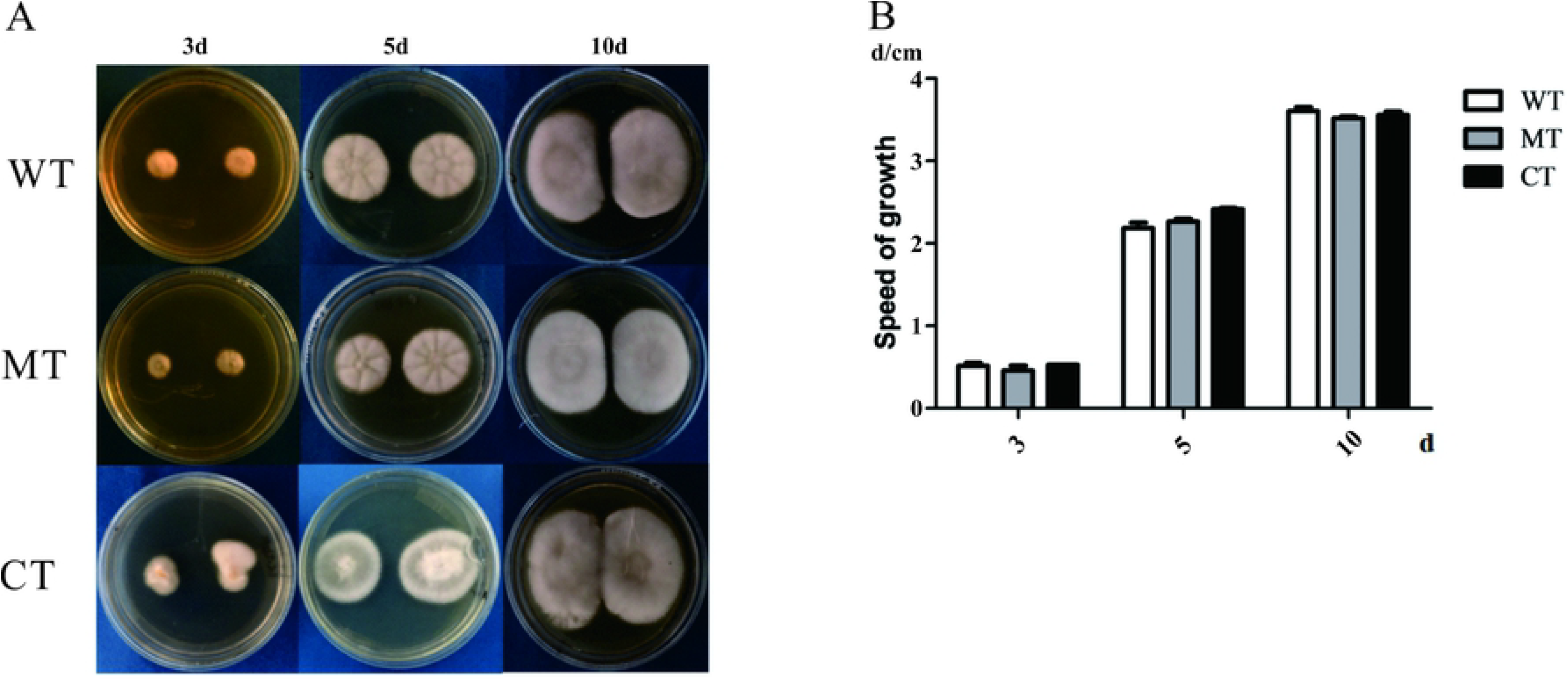
Phenotypic observation and growth rate determination of MT, CT and WT. (A-B) The MT, CT and WT of the same size were inoculated into the same position of the SDA medium at 28 °C for 3d, 5d and 10d, after which they were measured for diameter and photographed.

### Q Exactive Mass Spectrometer detection of SW in fermentation broth of MT, CT and WT

The same volume of MT, CT and WTfermentation broth was concentrated, and the concentration of SW was detected using a Q Exactive Mass Spectrometer. The peak time in the test samples and the SW standard was 4.4 (Fig.S8A, S8B and S8C). From the linear regression equation of mass concentration-peak area of SW, the content of SW in the fermentation broth of *M. anisopliae* was calculated to be 82.91 ± 15.92 μg/mg for WT (Fig.S8A), 56.42 ± 10.82 μg/mg for CT (Fig.S8C) and 5.71± 2.23 μg/mg for MT (Fig.S8B). The content of SW in the fermentation broth of the MT was significantly lower than that in the complemented strain and WT (Fig.6).

**Fig. 6.**
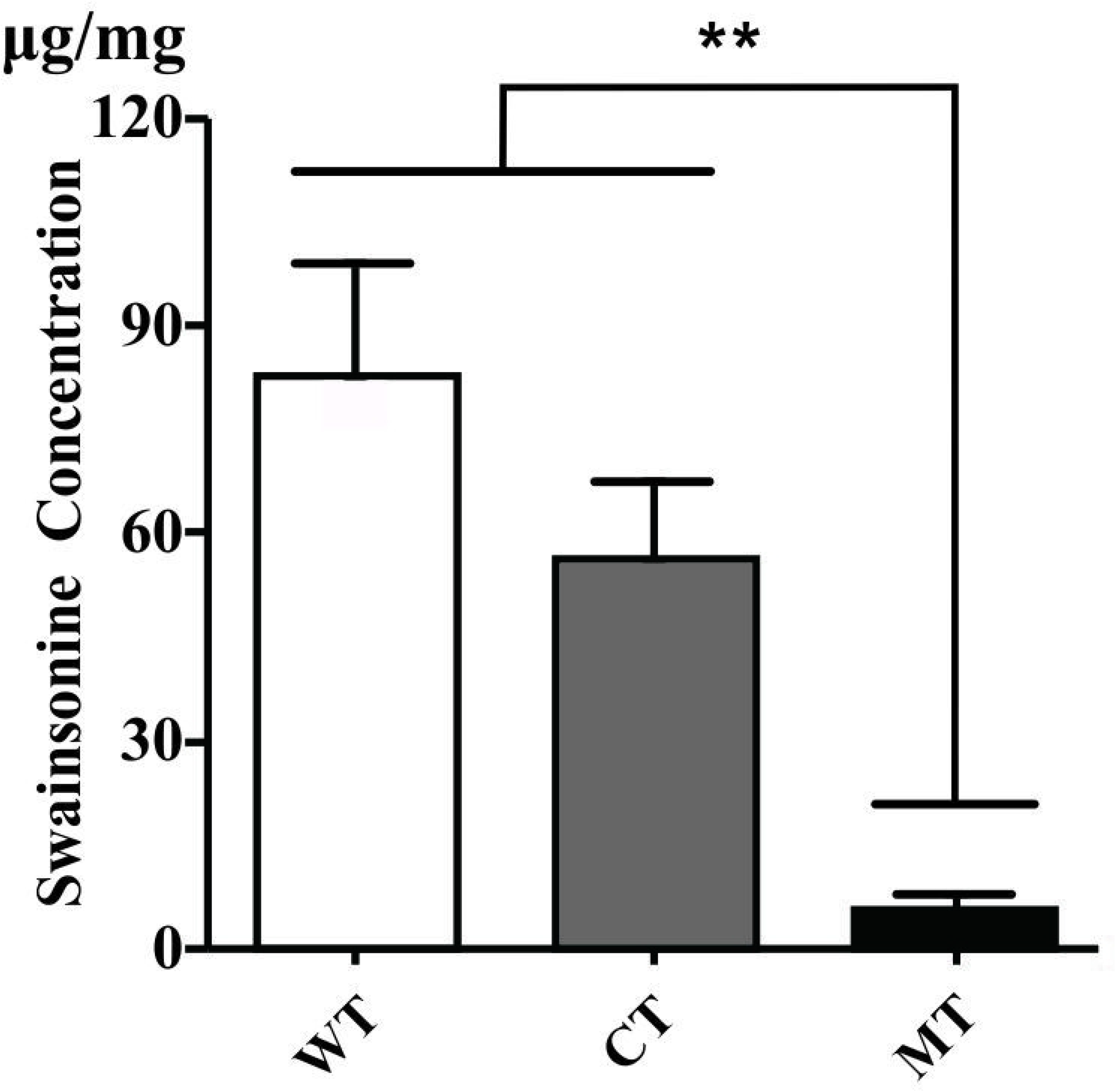
Q Exactive Mass Spectrometer detection of SW in fermentation broth of MT, CT and WT. The content of SW in the fermentation broth of the WT was significantly lower than that in CT and wild-type strain, and there was a significant difference. Data represented as mean ± SEM, n = 3, **P < 0.01.

## Discussion

While Cook et al. [17] demonstrated that the *swnK* gene is required for the SW biosynthesis of *M. robertsii*, no studies had determined which other genes are required for SW biosynthesis in *M. anisopliae*. We carried out a time course of SW in the fermentation broth of *M. anisopliae*, and compared the SWN gene expression for the same time course. The results showed that SW concentration was the highest at 3 d and the *swnR* gene was highly expressed at this time. This suggested that the *swnR* gene may play an important role in the SW biosynthesis pathway of *M. anisopliae*.

To further confirm the role of the *swnR* gene in the SW biosynthesis pathway of *M. anisopliae*, the *swnR* gene was knocked out using the benomyl resistance gene as a screening marker. The content of SW in MT was significantly reduced. Cook et al. [17] found that the swnK MT did not produce SW at all. However, in this study, after the *swnR* gene was knocked out, the content of SW was reduced, rather than completely absent. The cause of the low SW content may be due to the presence of a catalytic enzyme gene having the same function as the *swnR* gene. Since *swnN* is also a Rossman fold reductase, and is present in all SW-producing fungi, this could provide a similar activity and thus explain why SW was decreased rather than eliminated.

To demonstrate that the decrease in the content of SW was caused by the knock out of the *swnR* gene, the gene was complemented by inserting the wild-type *swnR* gene into the MT. The content of SW in the complement returned to normal levels, as predicted. Cook et al. [17] showed an increase in the content of SW after complementing the *swnK* gene, likely due to the high expression of the *swnK* gene in the complemented strain. We performed phenotypic observations and growth rate measurements on the MT, and found that it did not differ significantly from the wild-type strain. Therefore, the deletion of the *swnR* gene resulted in a decrease in the content of SW, but did not affect the radial growth of *M. anisopliae*. It is not known if sporulation or some other growth component was changed with the knockout.

The factors affecting the preparation of protoplasts and regeneration of filamentous fungi include the age of fungi, the choice of medium and enzymatic hydrolysate, enzymatic hydrolysis conditions and length of enzymatic hydrolysis [21, 22]. We found that *M. anisopliae* grown on SDA medium for 3d produced the most protoplasts. The cell wall composition of *M. anisopliae* is reported to be very complex [23], which might have favored the compound enzyme for the release of the *M. anisopliae* protoplasts This study showed that the protoplast release of *M. anisopliae* was the highest when digested with 1% snail enzyme, 1% cellulose, and 1% lysing enzymes for 3 h.

SW can cause neurotoxicity in grazing animals [24–26] and seriously threatens the production and development of animal husbandry [8, 9, 27]. SW also has significant anticancer and antitumor effects [28–30]. However, the limited source of SW, the difficulty of artificial synthesis, low extraction efficiency and high market prices have greatly limited the development of SW for anticancer and anti-tumor applications [14, 31, 32]. The production of SW by microbial fermentation, specifically by *M. anisopliae* or *S. leguminicola*[33] might be possible if the SWN pathway is better characterized.

## Conclusions

In this study, we demonstrated that the deletion of the *swnR* gene resulted in a decrease in swainsonine concentration in the fermentation broth of *M. anisopliae* and the recovery of swainsonine concentration to normal levels in the complemented strain. This suggests that the *swnR* gene plays an important role in the swainsonine biosynthesis pathway of *M. anisopliae* (Fig. 7). This study provides a preliminary research basis for the in-depth study of the swainsonine biosynthesis pathway and related catalytic enzyme genes.

**Fig. 7.**
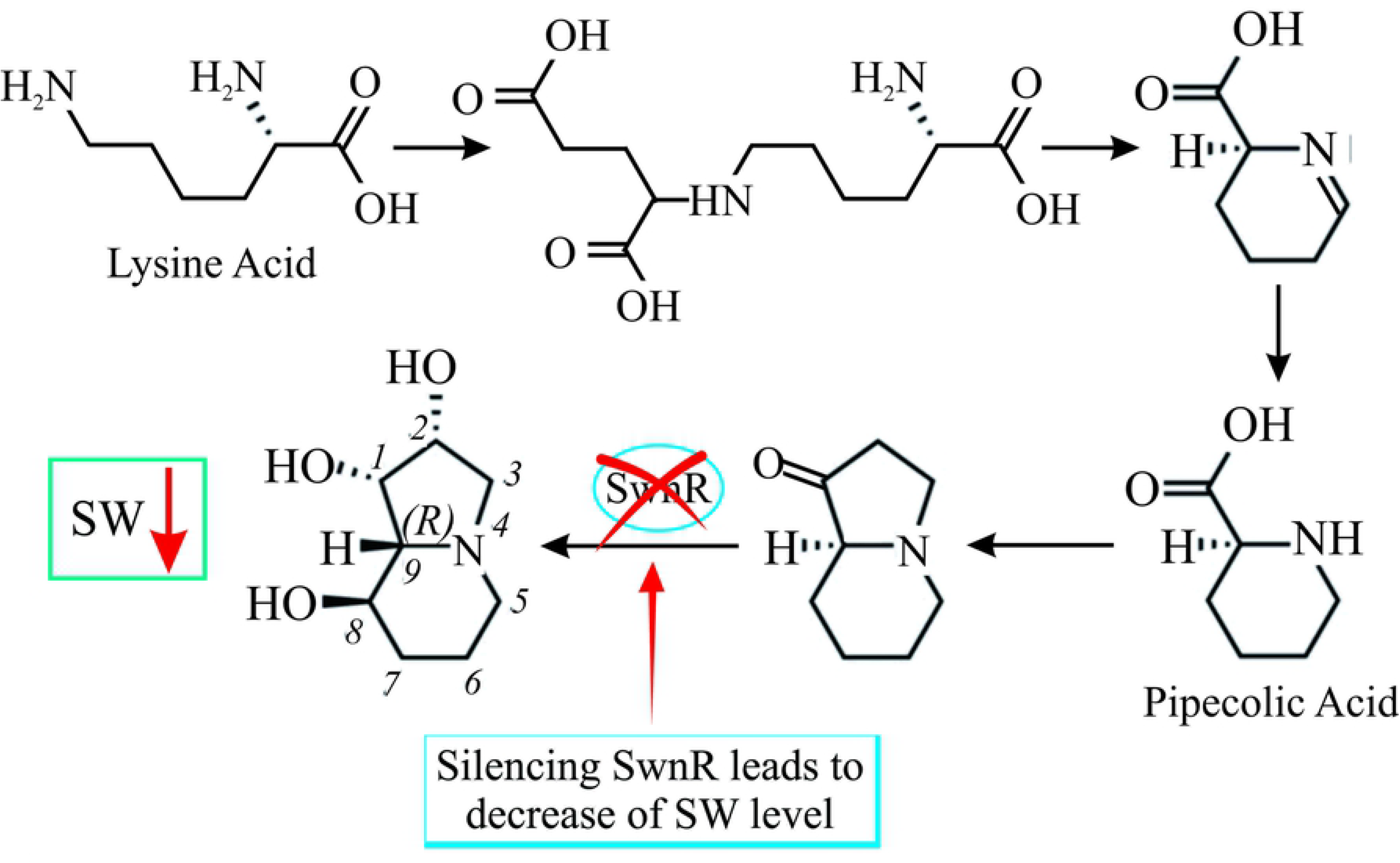
Decreased swainsonine content after knockout of SwnR gene by homologous recombination.

## Author Contributions

**Conceptualization:** Hao Lu, Baohai Wang, Baoyu Zhao, Qingyun Guo.

**Data curation:** Jingling Wang, Chonghui Mo.

**Formal analysis:** Lu Sun, Runjie Song, Yu Zhang, Yanli Zhu, Yiling Liu.

**Funding acquisition:** National Natural Science Foundation (No. 31201958), Key project of Natural Science Foundation of Shaanxi province (No. 2017JZ004), Science and Technology Special Fund Aid to Qinghai (2020-QY-210).

**Investigation:** Lu Sun, Runjie Song, Yu Zhang, Yanli Zhu, Yiling Liu.

**Methodology:** Lu Sun, Runjie Song.

**Project administration:** Hao Lu.

**Resources:** Hao Lu, Baoyu Zhao.

**Supervision:** Jinglong Wang, Qingyun Guo, Chonghui Mo, Baohai Wang, Baoyu Zhao, Hao Lu.

**Validation:** Lu Sun, RunJie Song.

**Visualization:** Lu Sun, RunJie Song.

**Writing-original draft:** Lu Sun, RunJie Song.

**Writing-review & editing:** Lu Sun, Runjie Song, Jinglong Wang, Yiling Liu, Yu Zhang, Yanli Zhu, Qingyun Guo, Chonghui Mo, Baohai Wang, Baoyu Zhao, Hao Lu.

